# Host plant affects the larval gut microbial communities of the generalist herbivores *Helicoverpa armigera* and *Spodoptera frugiperda*

**DOI:** 10.1101/2023.03.08.531690

**Authors:** Pedro A. P. Rodrigues, Nathalia Cavichiolli de Oliveira, Celso Omoto, Thomas Girke, Fernando L. Cônsoli

## Abstract

Investigations on symbiotic associations between Lepidoptera and bacteria have been inconclusive as to whether these microorganisms represent transient or persistent associations. One reason for this is that most studies have sampled the microbiota from only a few species and frequently from laboratory-reared animals. We sequenced the 16S rRNA gene to profile the gut microbiome of the field collected, agricultural pests *Helicoverpa armigera* and *Spodoptera frugiperda*. We found that variation in the structure of bacterial communities is in great part associated with their host diet. Moreover, *Enterococcus*, among a few other taxa, is consistently present in all samples regardless of the plant hosts were collected. The larval gut bacteria composition matches previously published data in the same or related species, indicating that associations with bacteria taxa such as *Enterococcus* are persistent despite geographical variation and sampling techniques. Based on our phylogenetic analysis it is not clear, however, if *Enterococci* associated with Lepidoptera are necessarily different from environmental bacteria. We conclude that to fully understand the nature of Lepidoptera - gut bacteria associations it will be necessary to conduct larger sampling efforts and combine them with identification methods of bacterial strains that go beyond screening of the 16S rRNA gene.

## Introduction

The evolution of herbivory in animals is often accompanied by the development of a beneficial association with microorganisms capable of making available nutrients that the host would not acquire otherwise (Ley et al., 2008; Hansen & Moran, 2013). Besides digesting plant tissues and synthesizing nutrients, there is growing evidence supporting the hypothesis that the gut microbiome facilitates an herbivorous lifestyle by detoxifying plant tissues (Kohl et al., 2014; Hammer & Bowers, 2015; Shikano et al., 2017; Itoh et al., 2018). Therefore, the herbivore gut microbiome is key to its host health, and its composition and structure are likely influenced by the type of plant consumed, its nutritional content, and the anti-herbivory toxins that it carries. A shift in diet in these animal host-microbe-plant interactions poses a challenge for both microbes and host. For instance, polyphagous moths may consume multiple plant species that vary in both nutritional composition and their anti-herbivory defense chemicals (Singer, 2008). Polyphagous moths harbor gut microbes that may be beneficial (Paniagua Voirol et al., 2018), but it is not yet well understood how these microorganisms are influenced by the characteristic wide diet breadth of their hosts.

Gut bacterial communities are often associated with digestive functions. Higher termites for instance rely on both protozoans and bacteria for digestion of cellulose and other plant fibers (Brune, 2014). The enzymes from metabolically diverse bacteria can help by either direct digestion or by the synthesis of vitamins and other nutrients that might otherwise be scarce in the host diet. In turtle ants (*Cephalotes*), for instance, gut bacteria are equipped with genes that can aid in the breakdown of nitrogenous waste and its conversion into essential amino acids (Hu et al., 2018). In addition to digestive functions, gut symbionts may also interact with other aspects of their host physiology. Of particular interest is the fact that some symbionts have been found to be capable of digesting chemicals associated with plant defenses against herbivory, as well as pesticides. As an example, *Candidatus* Ishikawaella capsulata, a gut bacterium found in kudzu bugs *Megacopta* spp. (Hemiptera: Plataspidae), has been found to be able to synthesize oxalate decarboxylase, which can help kudzu bugs to overcome oxalates produced as a defense barrier by the plants they eat (Nikoh et al., 2011). In the bean bug *Riptortus pedestris* (Hemiptera: Alydidae), bacteria in the genus *Burkholderia* have been shown to reduce the toxicity of fenitrothion to the host insect (Kikuchi et al., 2012). These results shed new light on how insects benefit from their symbionts and encourage a new perspective on the development of strategies for pest control.

These beneficial associations are often the result of co-evolution between a diet specialist and their symbiotic bacteria. Both *Megacopta* spp. and *R. pedestris* are examples of plant specialists, with a diet restricted to one or few related species of plants. But how is the ecology of microbiota in host species that are highly polyphagous? Many species in the moth family Noctuidae are highly polyphagous and also share great economic importance due to the damage they cause to a variety of crops worldwide. Bacteria found in a few species of Lepidoptera show metabolic functions that may be beneficial to their hosts, such as: (1) cellulolytic activity (Manfredi et al., 2015) and crystallization of alpha and beta carotenes (Shao et al., 2011), both found in bacterial cultures isolated from *Spodoptera;* (2) pesticide degradation reported in gut bacteria from laboratory-lines of *Plutella xylostella* (Ramya et al., 2016), as well as in laboratory and field-collected gut bacteria of both *Spodoptera frugiperda* (Almeida et al., 2017; Gomes et al., 2020) and *Helicoverpa armigera* (Madhusudhan et al., 2021); (3) and cases of changes in life history traits, such as laboratory-reared wax moth *Galleria mellonella* that were shown to live longer when inoculated with *Enterococcus* (Johnston & Rolff, 2015), a bacterium genus commonly found associated with Lepidoptera (Paniagua Voirol et al., 2018). *Enterococcus* from the gut of the fall armyworm also improved the performance of caterpillars, facilitating the use of a suboptimal diet (Chen et al., 2022).

Despite the importance of these findings, we highlight that most of these studies have been limited to laboratory-reared insects and their microbiota may not necessarily reflect the microbiota found in naturally occurring populations. For instance, *Enterococcus* occur in laboratory reared *Chloridea* moths, but not in field collected individuals (Staudacher et al., 2016). Therefore, one of the first steps to fully understand the interaction between highly polyphagous moths and their microbiota is to survey naturally occurring populations in different host plants. In the present study we sampled bacterial communities from the digestive tract of field collected *Spodoptera frugiperda* and *Helicoverpa armigera*, two of the most economically important moths, commonly found in a variety of crops worldwide. Our approach includes sequencing of the 16S rRNA gene to identify bacteria, and careful distinction between possible contaminants from likely resident bacteria, in addition to profiling of communities present in individuals fed different diets. With this combination we hope to contribute to a better understanding of the symbiotic relationships of gut bacteria and their Lepidoptera hosts.

## Results

The total number of raw sequences in samples of *S. frugiperda* was 2,094,934 and in *H. armigera* samples was 2,515,138. After quality trimming and filtering we were able to keep 239,882 sequences for *S. frugiperda* and 268,521 for *H. armigera*. The clustering of sequences into Operational Taxonomic Units (OTU) resulted in 202 OTUs for *H. armigera* and 427 for *S. frugiperda*. Most of these OTUs were present in low abundance. The top 98% of cumulative abundance of all samples (by insect host) is represented by just 34 OTUs for *H. armigera* and 18 OTUs for *S. frugiperda* (Fig. 1 A-B; Supplementary Tables S1 and S2).

**Fig. 1.**
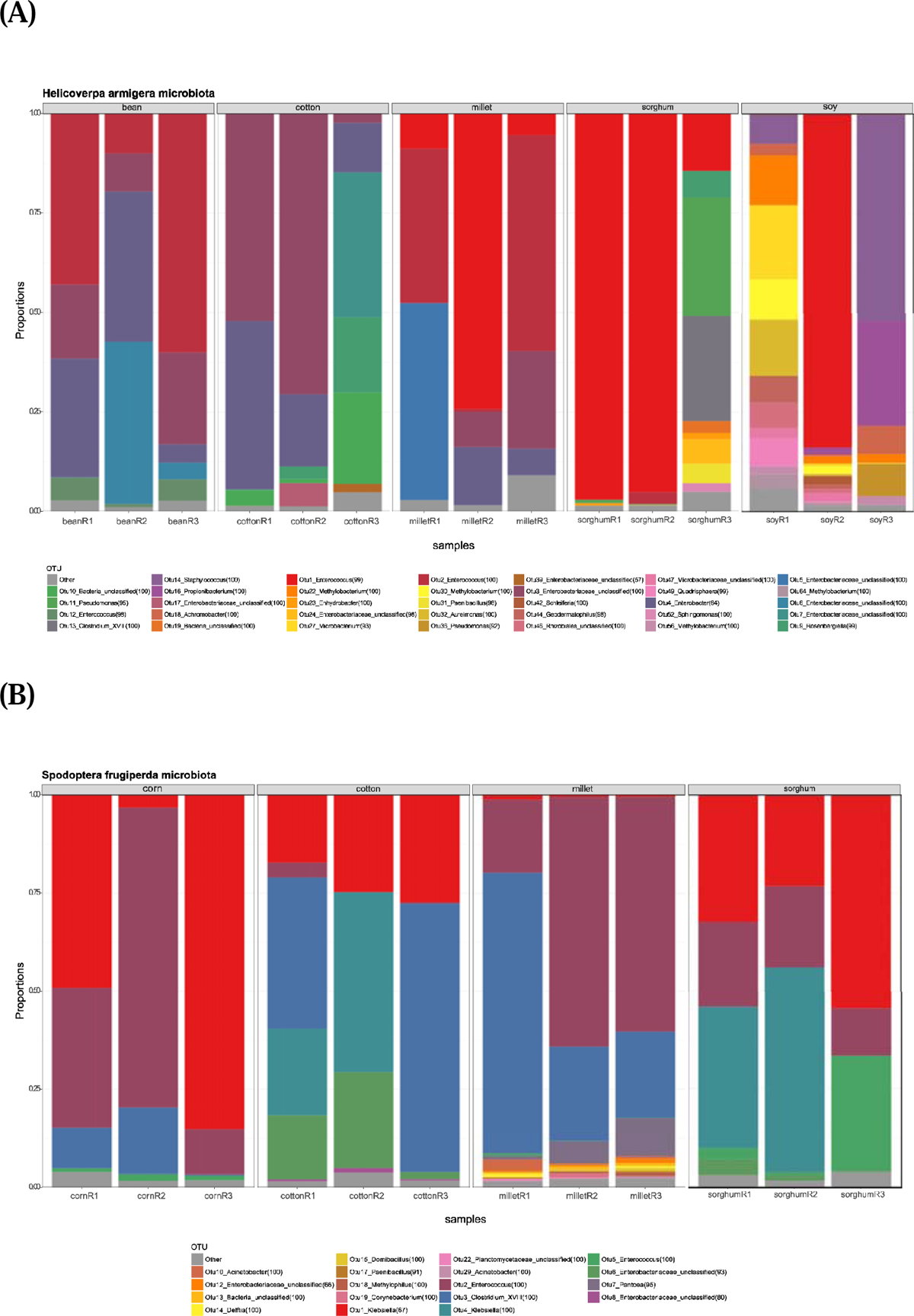
The gut microbiota of (A) *Helicoverpa armigera*, and (B) *Spodoptera frugiperda*. Each column represents a replicate within a diet group, where hosts were sampled. Each OTU is represented by a different color, and the number of reads associated with an OTU is represented by its relative proportion within the sample. “Other” are defined per diet group, and represent OTU that were found with relative abundance smaller than 2%.

We found that bacterial communities vary according to their host diet, for both *H. armigera* (PERMANOVAG, *alpha* = 0.5, *pseudo-F_4,14_* = 3.25, *p* = 0.001, *n* = 15) and *S. frugiperda* (PERMANOVAG, *alpha* = 0.5, *pseudo-F_3,11_* = 3.72, *p* = 0.006, *n* = 12). Despite the variation that we observed across replicates within diets (Fig. 1), the bacterial communities from larvae fed on monocots tended to be more similar to other monocots, and dicots to other dicots (Figure 2). An exception was found in the similar microbiotas of *H. armigera* larva collected in millet and in beans (Fig 2A): these samples shared OTU2 (*Enterococcus*), OTU3 (unclassified Enterobacteriaceae), and OTU4 (*Enterobacter*) as the most dominant bacteria. The variation in community composition was smallest for cotton samples and highest for soybean (Figs. 1A and 2A). Similarly, *Enterococcus* (OTU2) was also found to be dominant in *S. frugiperda*, particularly in the similar communities found in corn and sorghum. Insects fed on these two crops also shared *Klebsiella* (OTU1) as a dominant component (Fig. 1B). The variation within diet groups in *S. frugiperda* was observed to be generally smaller than the one observed in *H. armigera* (Figs. 1A-B, 2A-B). When considering bacteria taxa found in all samples OTU1 *Enterococcus* and OTU4 *Enterobacter* dominated the bacterial communities of *H. armigera*, while six OTUs dominated the bacterial communities in *S. frugiperda*: OTU1 *Klebsiella*, OTU2 *Enterococcus*, OTU3 *Clostridium*, OTU5 *Enterobacter*, and two other unclassified *Enterobacteriaceaea* (OTUs 6 and 8) (Table 1).

**Fig. 2.**
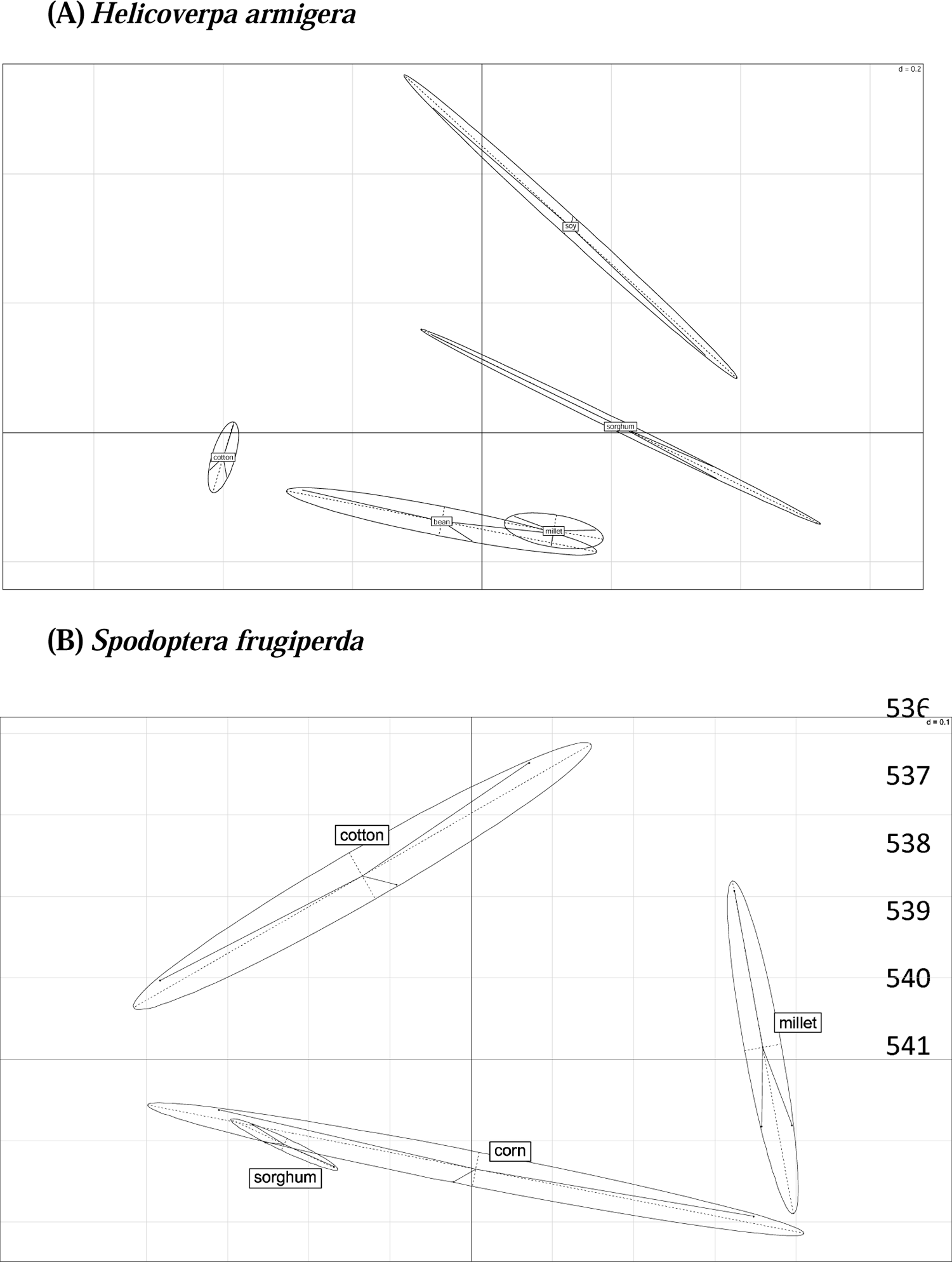
Principal components analysis. Grouping of samples according to dissimilarity Unifrac (alpha=0.5) distances, for (A) *Helicoverpa armigera* (k=2, GOF=0.59), and (B) *Spodoptera frugiperda* (k=2, GOF=0.75). The distance of each square side is given as a “d” value in each plot.

**Table 1.**
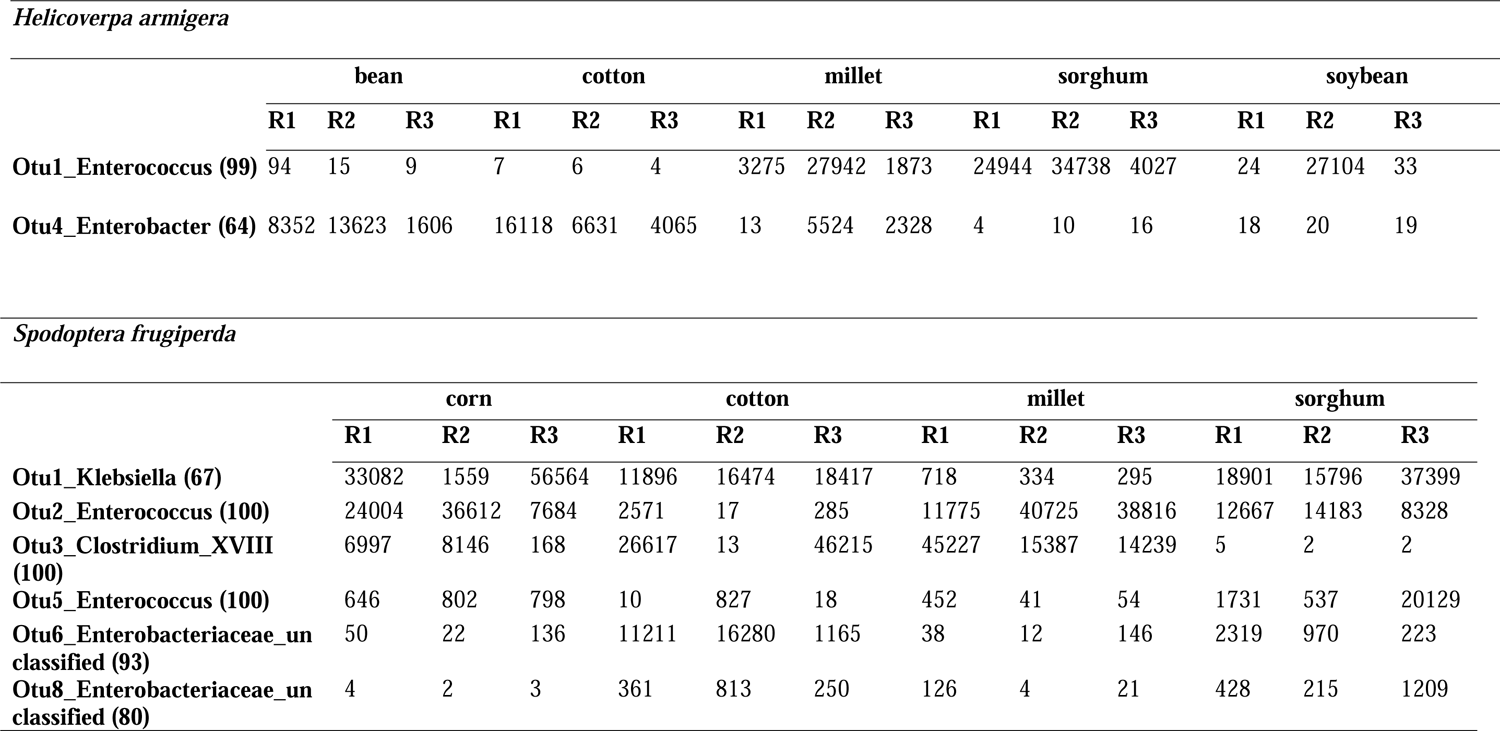
Frequency of OTU that are present in all samples across diet groups, for each host species. Top: *Helicoverpa armigera*; Bottom: Spodoptera frugiperda.

|  | bean |  |  | cotton |  |  | millet |  |  | sorghum |  |  | soybean |  |  |
| --- | --- | --- | --- | --- | --- | --- | --- | --- | --- | --- | --- | --- | --- | --- | --- |
|  | R1 | R2 | R3 | R1 | R2 | R3 | R1 | R2 | R3 | R1 | R2 | R3 | R1 | R2 | R3 |
| Otu1_Enterococcus (99) | 94 | 15 | 9 | 7 | 6 | 4 | 3275 | 27942 | 1873 | 24944 | 34738 | 4027 | 24 | 27104 | 33 |
| Otu4_Enterobacter (64) | 8352 | 13623 | 1606 | 16118 | 6631 | 4065 | 13 | 5524 | 2328 | 4 | 10 | 16 | 18 | 20 | 19 |

*Spodoptera frugiperda*
|  | corn |  |  | cotton |  |  | millet |  |  | sorghum |  |  |
| --- | --- | --- | --- | --- | --- | --- | --- | --- | --- | --- | --- | --- |
|  | R1 | R2 | R3 | R1 | R2 | R3 | R1 | R2 | R3 | R1 | R2 | R3 |
| Otu1_Klebsiella (67) | 33082 | 1559 | 56564 | 11896 | 16474 | 18417 | 718 | 334 | 295 | 18901 | 15796 | 37399 |
| Otu2_Enterococcus (100) | 24004 | 36612 | 7684 | 2571 | 17 | 285 | 11775 | 40725 | 38816 | 12667 | 14183 | 8328 |
| Otu3_Clostridium_XVIII (100) | 6997 | 8146 | 168 | 26617 | 13 | 46215 | 45227 | 15387 | 14239 | 5 | 2 | 2 |
| Otu5_Enterococcus (100) | 646 | 802 | 798 | 10 | 827 | 18 | 452 | 41 | 54 | 1731 | 537 | 20129 |
| Otu6_Enterobacteriaceae_unclassified (93) | 50 | 22 | 136 | 11211 | 16280 | 1165 | 38 | 12 | 146 | 2319 | 970 | 223 |
| Otu8_Enterobacteriaceae_unclassified (80) | 4 | 2 | 3 | 361 | 813 | 250 | 126 | 4 | 21 | 428 | 215 | 1209 |

To build a 16S rRNA phylogenetic tree we processed 305 sequences downloaded from NCBI, in addition to our outside group sequence, *Lactiplantibacillus plantarum* (NR_042394.1). After alignment and quality trimming, 219 sequences were kept. These sequences were grouped in 12 clusters of 99% similarity (or a dissimilarity distance of 0.01, also known as exact sequence variants, or ESV, see Supplementary Table S3). We found that while *Enterococcus* ESVs associated to Lepidoptera were very similar to each other, they also included sequences associated to a variety of other environments, including humans, agriculture, waste, water, and soil (Supplementary Table S3). To gain further insight, we picked the most abundant OTUs identified as *Enterococcus* across our samples (Table 1) and aligned them against the 12 *Enterococcus* ESV sequences. When aligned to OTU1 from *H. armigera* and OTU2 from *S. frugiperda* – which are identical to each other – we found that two *E. mundtii* ESVs contain sequences that share 100% identity to the region sequenced. OTU2 from *H. armigera* and OTU5 from *S. frugiperda* were found to share 100% identity to sequences in the cluster ESV 02 corresponding to *E. casseliflavus*. In addition, OTU2 from *H. armigera* also shared 100% identity to another *E. casseliflavus* cluster, ESV 03 (Supplementary Table S4).

## Discussion

The gut microbiota of *S. frugiperda* and *H. armigera* are relatively simple, and most of the sequences belong to a few, dominant OTUs. Crop origin (diet) explained most of the variation observed among gut bacterial communities (Figs. 1 and 2), consistent with findings in other animals, including other Lepidoptera, where diet was also found to be associated to changes in gut microbiota (Hu et al., 2014; Flint et al., 2015; Matteo et al., 2016; Wang et al., 2020). Despite changes with diet, a few abundant OTUs were found to be dominant regardless of diet or sample size (Table 1).

A composition of a few but highly abundant taxa is consistent with how gut bacterial communities are structured in other insects (Anderson et al. 2016; Lanan et al. 2016). According to a review on Lepidoptera microbiota, *Enterococcus* has been reported in approximately 70% of the species studied, and *Klebsiella* in about 35% (Paniagua-Voirol et al., 2018). Our results show that these two genera were also found within the core microbiota for both *H. armigera* and *S. frugiperda.* In addition, we found abundant unclassified OTUs in the family *Enterobacteriaceae*, which have also been reported to be associated with more than 80% of species of Lepidoptera studied to date (Paniagua Voirol et al., 2018). It is important to highlight that although we identified OTUs as *Enterococcus* with bootstrap values near 100%, OTUs identified as *Klebsiella* likely represent a different genus, as their bootstrap value was relatively low, 67%. This uncertainty about *Klebsiella* and the ubiquitous but unclassified *Enterobacteriaceae* both illustrate the necessity to better characterize microorganisms found in association with Lepidoptera, to allow for more meaningful inferences about their similarity to environmental microorganisms and across different hosts.

### Symbionts or transient bacteria?

One of the criteria to define a bacterium as a symbiont is to distinguish between species that are just transiting the digestive tract along with the ingestion of food from actual residents that establish a more permanent association with their host. Are the community composition changes observed in *H. armigera* and *S. frugiperda* just a product of ingestion of environmental bacteria? In a survey of gut microbes in *H. armigera* from India it was reported that the caterpillars’ gut communities correlated with the bacterial composition found in food that they ingested, and the similarity between gut and food bacterial communities ranged from 73% to 93% (Gayatri Priya et al., 2012), suggesting that most gut bacteria are transient associates. In another study with Lycaenidae butterflies, it was found that most of the bacteria found in the digestive tract of caterpillars were also found in the soil and plant surfaces (Whitaker et al., 2016). Nevertheless, in this latter study, the authors reported that some variation could not be explained by diet, even when comparing carnivorous with herbivorous species, concluding that these results provide evidence that common environmental bacteria may be ingested with diet, but these microorganisms could still be relevant to their host’s biology (Whitaker et al., 2016). In fact, this is the case for other insects, such as in the case of the insecticide-degrading *Burkholderia*-bean bug symbiosis, in which their hosts must acquire this bacterium from the environment at every generation (Kikuchi et al., 2012). It has also been shown that even ingested bacteria can colonize and survive throughout development and generations of Lepidoptera. When GFP-tagged *Enterococcus mundtii* – a bacterium consistently found in *Spodoptera* – is fed to 3^rd^ instars, this bacterium not only survived the physical-chemical conditions of the gut, but also proliferated during developmental stages, and were detected in pupae, adults and even inside oocytes and first instars of the second generation (Teh et al., 2016). The persistence of a bacterium acquired through ingestion suggests that environmentally acquired bacteria can establish associations with Lepidoptera that last through developmental stages and generations. Furthermore, *Enterococcus* is frequently reported as a core member of the *S. frugiperda* gut microbiota, by different research groups and geography (Ugwu et al., 2020; Higuita Palacio et al., 2021; Oliveira et al., 2022). Altogether, these evidences support the hypothesis that Lepidoptera hosts are capable of developing associations with environmental bacteria, but only with more sampling and experimentation it will be possible to determine whether these microbes become resident and beneficial to their hosts.

To gain further insight on the nature of these associations we downloaded 16S rRNA sequences identified as *E. mundtii* and *E. casseliflavus* from NCBI, to determine whether sequences associated to Lepidoptera were more similar to each other than sequences from other environments. Our results showed that *Enterococcus* associated to Lepidoptera formed a single ESV for *E. mundtii* and another single ESV for *E. casseliflavus* (Fig 3). Although this result seems to support the hypothesis of a specific host-microbe association, these same two ESVs also contained sequences that were associated to a variety of other environments, including water, humans, and other vertebrates and invertebrates. When we look only for similarities between these sequences and OTU1 (*Enterococcus mundtii* in *S frugiperda* and *H. armigera*) and OTU5 (*Enterococcus casseliflavus* in *S. frugiperda*), we find that the target region that we sequenced (V3-V4) is identical to sequences that were sampled in the silk moth (*Bombyx mori*) but also in sheep, humans, and soil (Supplementary Table S3).

**Fig. 3.**
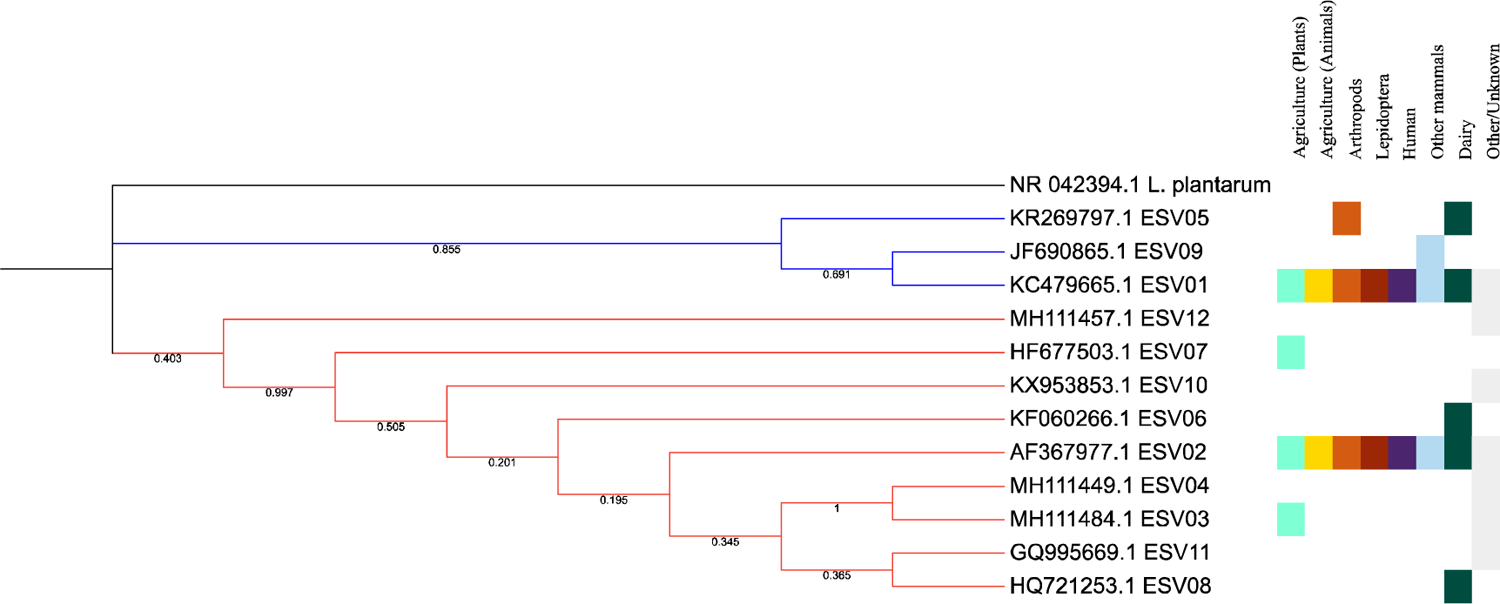
Maximum likelihood phylogenetic tree depicting 12 *Enterococcus mundtii* (blue branches) and *E. casseliflavus* (red branches) and their different environmental sources. *L. plantarum* is included as an outgroup. This topology represents the configuration with highest log likelihood, −3641.30. A Kimura-2 factors was selected as the model that best fit our data, with a discrete Gamma distribution to model evolutionary rate differences among sites (5 categories (+G, parameter = 0.2671)). Sequences were aligned using the Muscle algorithm and tree was built using MEGA11.

But are these sequences necessarily the same organism? The insecticide-degrading *Burkholderia insecticola* was recently described as a new species (Takeshita et al., 2018), nevertheless its 16S rRNA shows 100% identity to another species, *Burkholderia peredens* LMG 29314T (Peeters et al., 2016; Takeshita et al., 2018). The association *R. pedestris*-*B. insecticola* illustrates that the taxonomic resolution obtained by the 16S rRNA gene is limited and might become a confounding factor when trying to distinguish between possibly symbiotic bacteria from ubiquitous microorganisms such as *E. mundtii* reported in a variety of environments (Supplementary Table S3). We found that *H. armigera* OTU2, corresponding to *E. casseliflavus*, was identical to regions of sequences contained by both ESV 02 and ESV 03. Although ESV 02 includes sequences isolated from Lepidoptera, none of the sequences in ESV 03 was sampled from Lepidoptera or any other arthropod. Therefore, similarly to the *R. pedestris*-*B. insecticola* system, the identification of specificity between *Enterococcus* and *S. frugiperda* and *H. armigera* cannot be supported or rejected solely on information provided by 16S rRNA sequences.

We cannot rule out that *Enterococci* found in association with *H. armigera* and *S. frugiperda* may represent distinct species of bacteria, despite sharing up to 100% 16S rRNA identity with several other *Enterococci* (Supplementary Table S4). Future surveys and experiments should adopt additional approaches to distinguish from diet- or environment-associated bacteria from gut-associates, such as a shotgun-metagenome sequencing of the bacterium genome, or the development of a MLST or TRFLP system.

## Conclusion

In this study we investigated the gut community composition from two of the most economically important agricultural pests worldwide: *H. armigera* and *S. frugiperda*. We found that although some community members are ubiquitous to all samples, there was significant variation explained by type of diet hosts were consuming at the time they were collected. The current controversy regarding the status of symbionts for microbes associated with Lepidoptera is mostly derived from a paucity of studies on this subject. There are surprisingly few studies that have focused on investigating the role of microbes found in the gut lumen of herbivorous insects (Hansen & Moran, 2013). Finally, we conclude that the 16S rRNA DNA region may be insufficient for testing the hypothesis of a species-specific associations between *Enterococcus* and Lepidoptera. Different bacteria may share similar or identical sequences, and in the case of sequences identified as *E. mundtii* or as *E. casseliflavus,* we found identical 16S rRNA sequences originating from a diversity of environments and geographical regions, that are likely representatives of different strains/species.

### Experimental procedures

#### Samples

Larval stages of both *S. frugiperda* and *H. armigera* were collected in cotton, millet and sorghum plantations. *Spodoptera frugiperda* was also sampled from corn fields, and additional *H. armigera* samples were collected in beans and soybean fields. All collections were done in a farm landscape located in the state of Bahia, Brazil (12 05’30.84”S; 45 48’18.00”W). Samples were immediately preserved in RNAlater, and transported to the laboratory in São Paulo State, Brazil, for storage at −80°C until dissections and RNA extractions were done.

Larva had their head capsules measured, and fifteen 5^th^ instar larvae for each crop and insect species were randomly selected for dissection. The gut was dissected out and the midgut removed for the collection of the endoperitrophic content by pulling the peritrophic matrix out with fine forceps. Dissections were carefully done in sterile conditions to avoid external contamination and from sample-to-sample contamination. The peritrophic matrix containing the food content was pooled in groups of five and kept in RNAlater at −80°C. Three biological samples (1 sample = 5 pooled guts) were prepared for each treatment (insect species/host plant), and used for sequencing.

Samples were homogenized prior to DNA extraction, and for each sample an aliquot volume corresponding to 1.25 gut content equivalent was washed with a cold PBS solution for the elimination of the RNAlater solution through centrifugation (15 min, 12,000 *g*, 4°C). The top clear liquid layer was discarded, and the remaining volume subjected to DNA extraction. Samples obtained from larvae collected in cotton were subjected to an alternative DNA extraction protocol (Japelaghi et al., 2011) due to the high polyphenol content associated with them. All other samples were extracted as described below.

### DNA extraction and sequencing

After RNAlater removal, samples were incubated overnight in 750 μ of phenol:chloroform:isoamyl alcohol (PCI, 25:24:1) was added, and the samples were mixed rapidly by inversion for 2 min. Samples were centrifuged (8,000 g x 10 min), and the aqueous layer was collected and re-extracted twice with a similar volume of PCI. Next, chloroform was added at 1:1 ratio, the sample was shaken vigorously for 15 s and centrifuged as before. The obtained pellets were resuspended in 1/10 volume of sodium acetate (3M, pH 5.2) and 1 volume of ethanol (95%), incubated for 40 min at −80°C, and centrifuged (16,000 *g* x 30 min x 4°C). The pellet was washed twice with 1 mL of 85% ice-cold ethanol, and the pellet obtained was allowed to dry for 5 min in a SpeedVac at 65°C. The pellet was resuspended in ultrapure water and kept at −80°C. Libraries were prepared following the Illumina protocol for 16S metagenomics experiments. Briefly, the V3-V4 region of the 16S rRNA gene was PCR-amplified under the following conditions: the reaction was programmed at 95°C for 3 min (1 cycle), followed by 25 cycles at 95°C for 30 s, 55°C for 30s and 72°C for 30 s, with a final extension (1 cycle) at 72°C for 5 min. The amplicons were submitted to the molecular core center facility at ESALQ/USP (Brazil), to be sequenced in the Illumina MiSeq platform, using TruSeq chemistry for paired-end sequencing 2×250bp.

### Bioinformatics

Quality-trimming and OTU-picking was performed in the software Mothur (Schloss et al., 2009), following their Standard Operation Procedure for MiSeq data (Kozich et al., 2013), and downstream data analysis was done in R (R Core Team, 2013). Briefly, we quality trimmed sequences and discarded any sequence with an insert size longer than the region sequenced (i.e., minimal length of 445bp and maximum of 471bp), or that aligned outside of the region sequenced (using *E. coli* as a reference, positions start=6428, end=23440). Next, we removed sequences with more than 8 homopolymers, chimeric sequences and sequences that classified to phyla other than Bacteria, or that were not classified at the phylum level. Sequences were clustered into Operational Taxonomical Units (OTU) using the OptiClust algorithm, that has recently been shown to be one of the best to reduce the number of false positives and negatives (Westcott & Schloss, 2017). An OTU can be considered as a more conservative definition of species, as most microorganisms have not been described, and the concept of species is often controversial for Bacteria. The variation in community composition was investigated in R, with samples rarefied to the smallest sample count. For each diet (i.e., crop origin), samples were filtered as following: (1) all singletons were excluded, and (2) if an OTU was not present in 2 out of 3 replicates, it was deemed as a contaminant and further excluded from the analysis. We defined the “core” microbiota as the community of bacteria that has passed the two filters described earlier, and that were within the top 98% of all sequence counts. To find out if diet explained the variation observed across samples, we first calculated variable Unifrac distances (weight = 0.5) and used these distances to build a Principal Components Analysis plot. We further investigated the association of diet with microbial community composition by performing an analysis of variance with permutation, using both UniFrac distances and a maximum likelihood UPGMA tree with representative sequences of each OTU (PERMANOVAG, Chen et al. 2012).

To gain insight on the nature of bacteria associated with *S. frugiperda* and *H. armigera* (symbiont or transient), we identified which groups of bacteria were found in all samples regardless of sample source but selected from the groups of bacteria that passed the filters described earlier. In this survey we found that OTUs identified as *Enterococcus mundtii* and *Enterococcus casseliflavus* were found in all samples, which are taxa commonly reported for Lepidoptera. To further investigate these associations, we built a phylogenetic tree containing representative sequences of *E. mundtii* and *E. casseliflavus* to verify if the *Enterococci* associated with Lepidoptera share more similarities to each other than to *Enterococci* associated with other organisms. To build this tree we first downloaded 16S rRNA sequences that included regions of the 16S rRNA gene beyond the ones we sequenced (V3-V4). We searched the NCBI nucleotide database for 16S rRNA sequences identified as *E. mundtii* and *E. casseliflavus* and downloaded them if their length was between 1400-1600 base pairs. We also downloaded a 16S rRNA sequence for *Lactiplantibacillus plantarum* (NR_042394.1) that we defined as our outgroup. In Mothur, all *Enterococcus* sequences were aligned against the Silva database (v138) (Quast et al., 2013; Glöckner et al. 2017). Next, we used *screen.seqs* to select sequences with a minimum length of 1000 bp, starting at position 1046, with zero ambiguities, a maximum of 8 homopolymers, and optimizing the end position so that at least 90% of sequences are kept in the analysis. Next, we used *unique.seqs* to dereplicate sequences, and *filter.seqs* to eliminate columns of gaps or periods common to all sequences. Next, we calculate distances (“*dist.seqs*”, cutoff=0.03), clustered sequences (“*cluster*”, method=average) and selected representative sequences for each cluster at distance 0.01 (also known as exact sequence variants, ESVs). Using these representative sequences, we built a maximum likelihood phylogenetic tree with 1000 bootstrap replications, using the phylogenetic tree model Kimura 2-parameter with gamma distribution, selected in MEGA 11 as the best fit for our data. To manually add information on sample source and bacterial species, we used the interactive tree of life iTol online platform (http://itol.embl.de/) (Letunic & Bork, 2016).

The following R packages were used for data processing, statistical and phylogenetic analysis, as well as visualization: *vegan* (Oksanen et al., 2017), *ggplot2* (Wickham, 2009), *RColorBrewer* (Neuwirth, 2014), *GUniFrac* (Chen et al., 2012), *ade, ape* (Paradis et al., 2004), *phangorn* (Schliep, 2011), and *dplyr* (Wickham et al., 2015).

## Data availability

Raw sequencing data will be made available through NCBI upon acceptance for publication. Clustered sequences and filtered data will be available through the online data sharing platform FigShare. Finally, all steps used in data analysis were compiled as a single R script, available at our GitHub page [to be made available upon acceptance for publication].

## Acknowledgements

This project was funded by Conselho Nacional de Desenvolvimento Científico e Tecnológico (CNPq) (process 403851/2013 to CO; process 462140/2014-8 to FLC); São Paulo State Research Foundation (FAPESP) (process 2014/16609-7 to FLC; and 2016/11127-0 and 2017/03415-8 for fellowships to PAPR, and 2017/24377-7 for a fellowship to NCO). Computations were performed using the computer clusters and data storage resources of the UCR’s HPCC, which were funded by grants from NSF (MRI-1429826) and NIH (1S10OD016290-01A1).

## Author Contributions

FLC and CO designed the study, obtained all samples and secured the funds for this project; NCO processed all biological samples for collecting the gut microbiota; PAPR obtained DNA samples and generated amplicons for sequencing; PAPR and TG processed and analyzed the data; PAPR and NCO wrote the initial draft; FLC edited the draft; All authors read and approved the final version of the manuscript.

## SUPPLEMENTAL MATERIAL

**Table S1.**
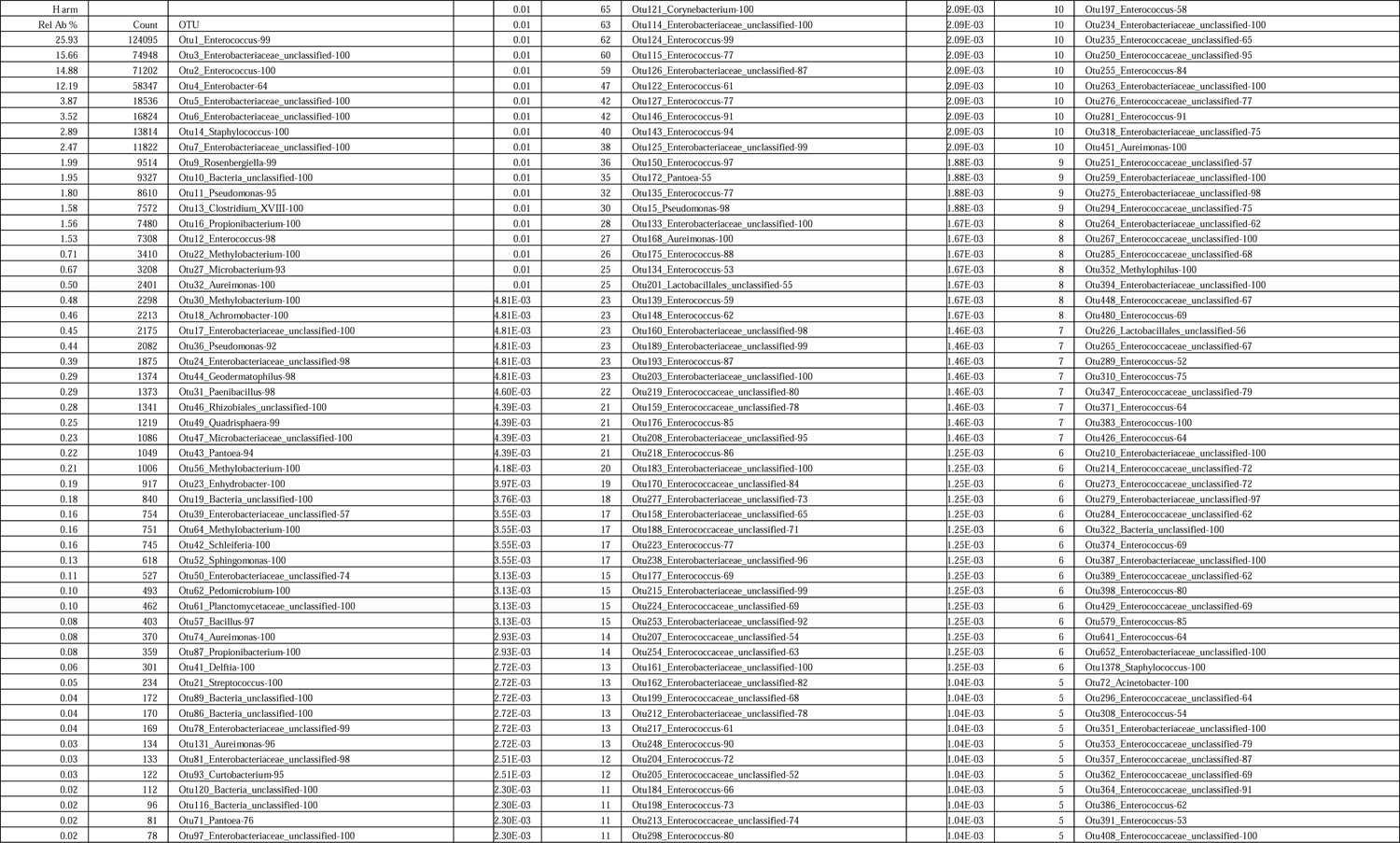
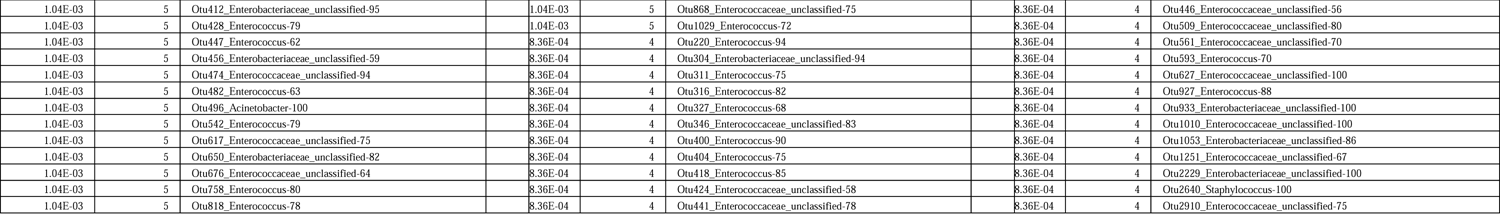
Total number of reads for each OTU identified in *Helicoverpa armigera*.

**Table S2.**
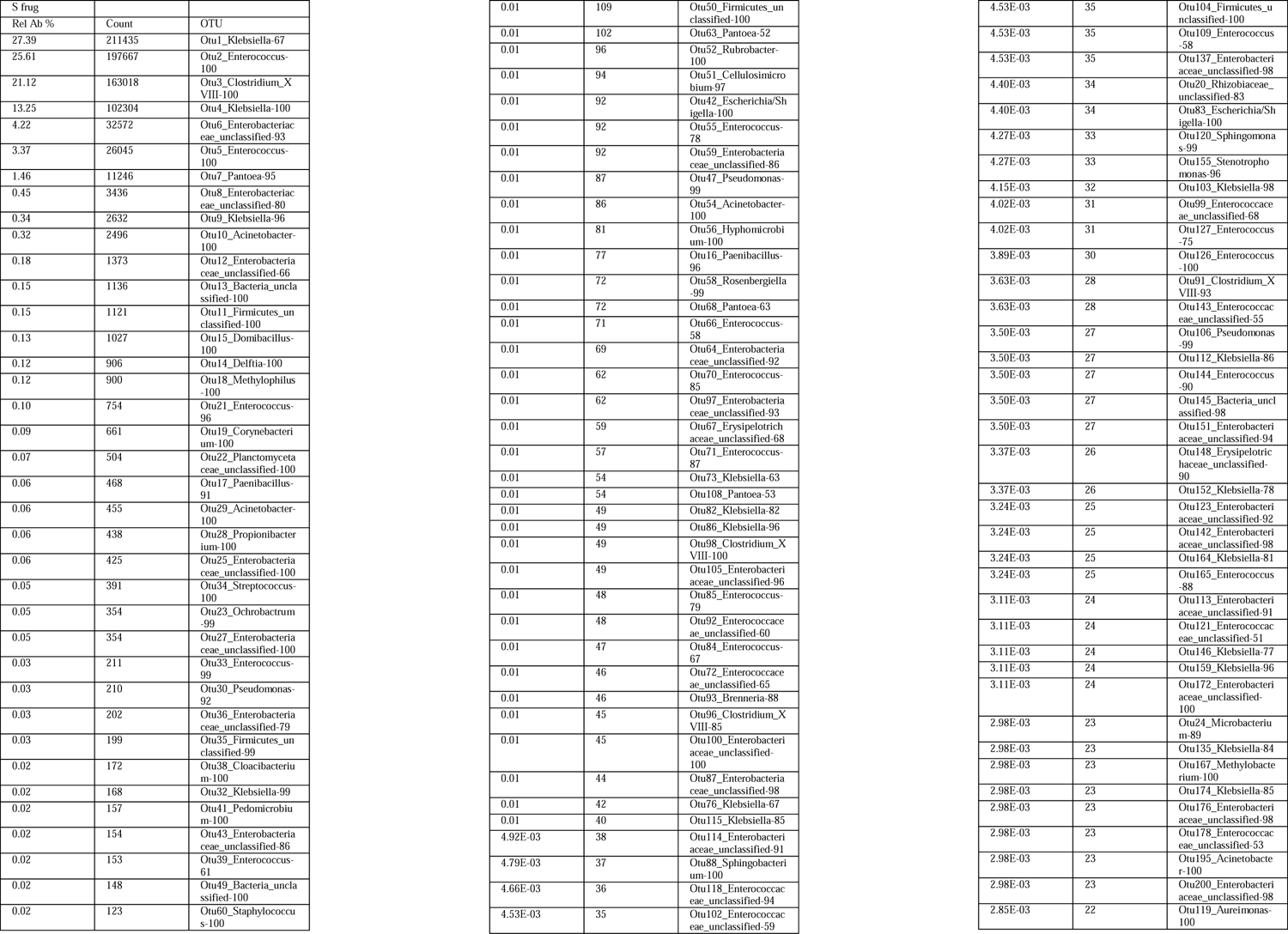
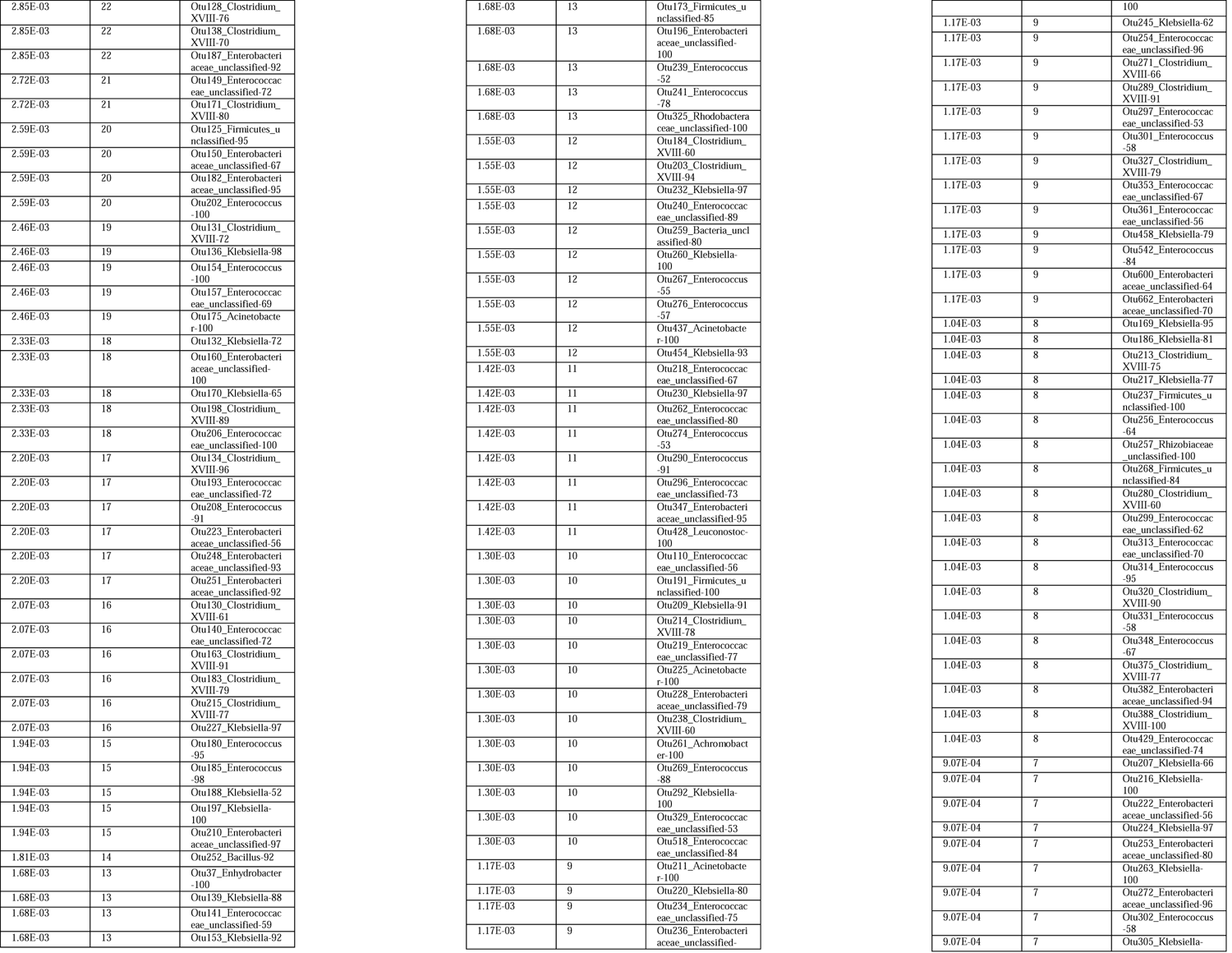
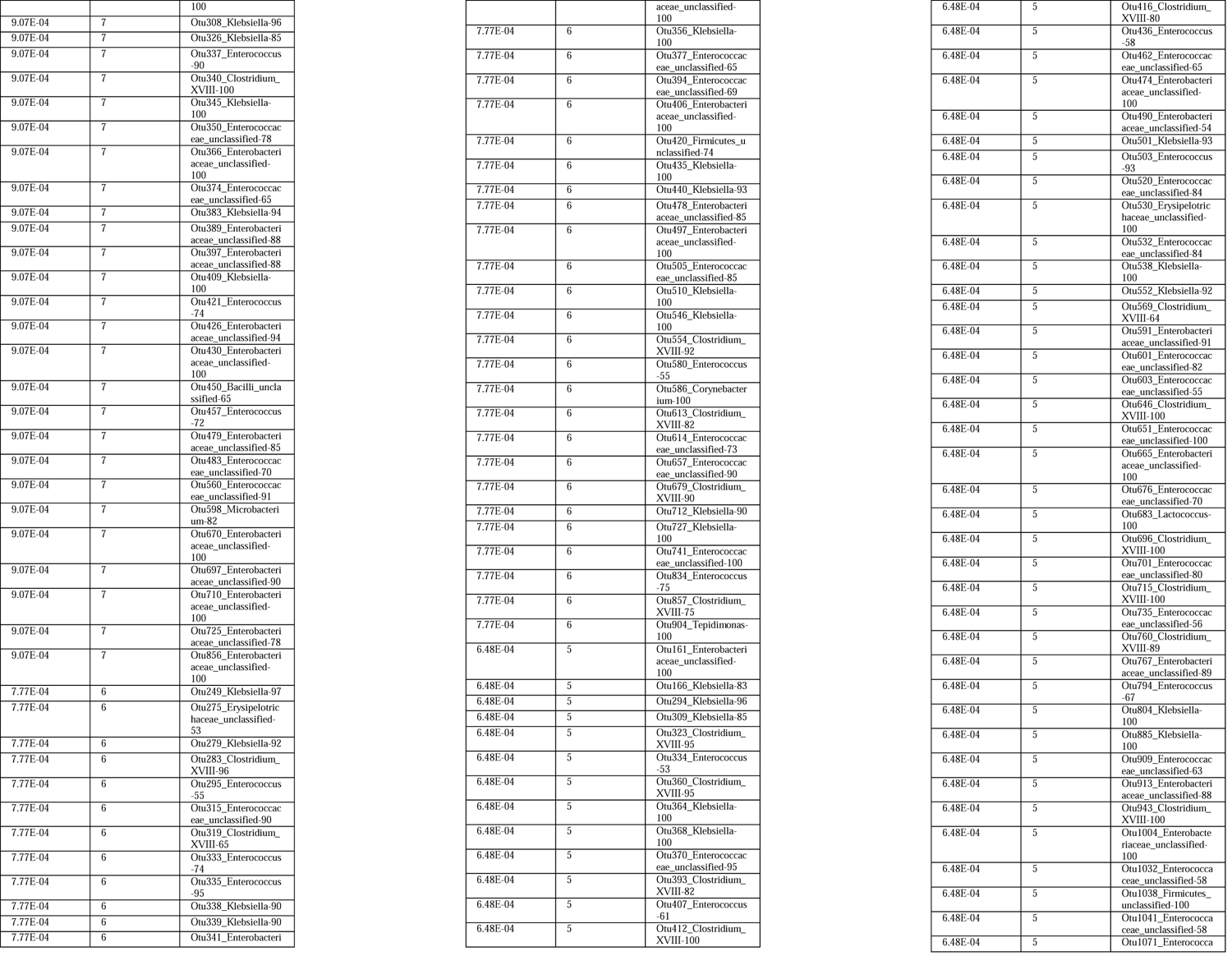
Total number of reads for each OTU identified in *Spodoptera frugiperda*.

**Table S3.**
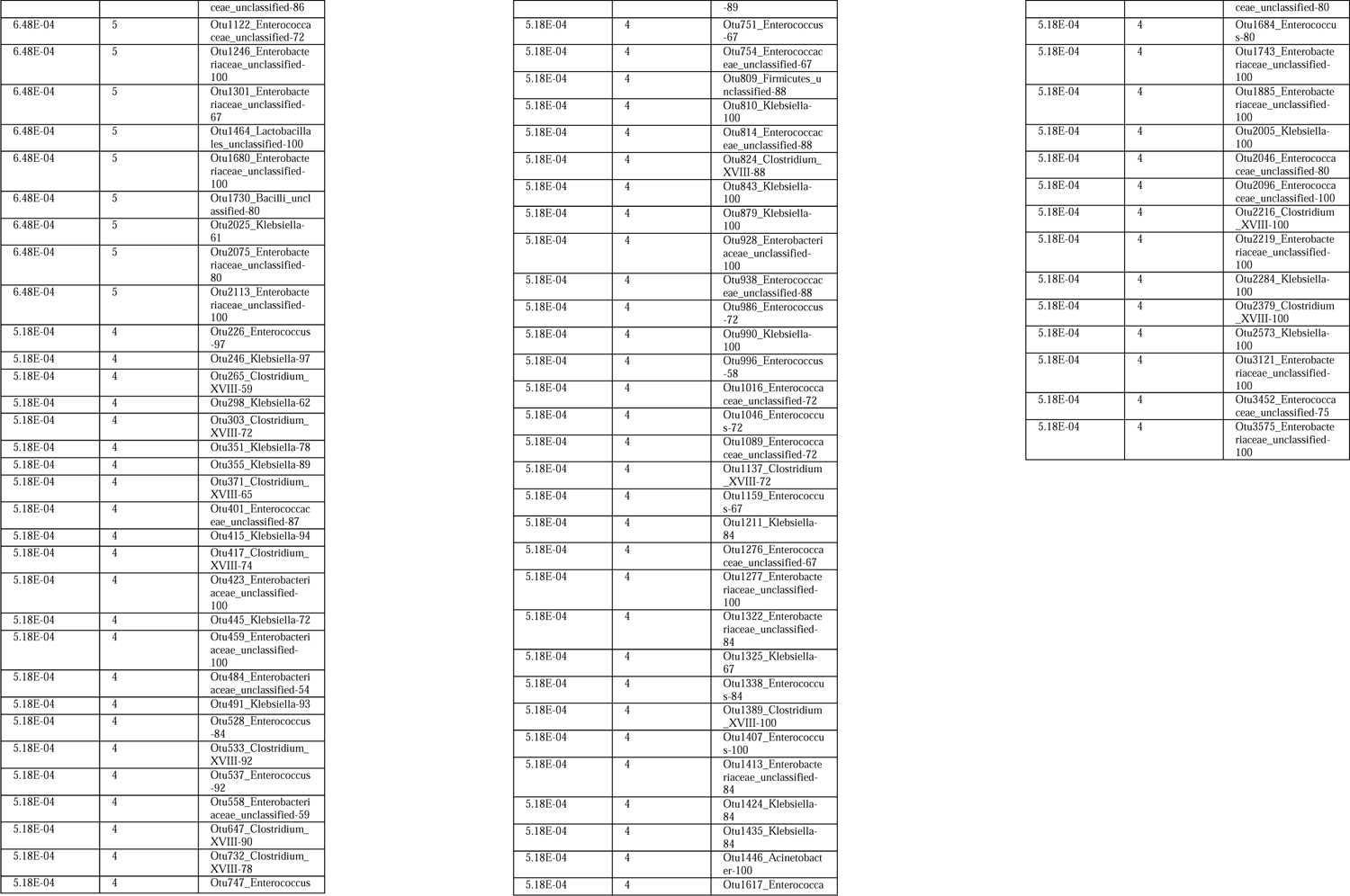
Metadata information for sequences used as input for the phylogenetic tree (Figure 3). Table provided as an Excel ® Spreadsheet (see attachment)

**Table S4.**
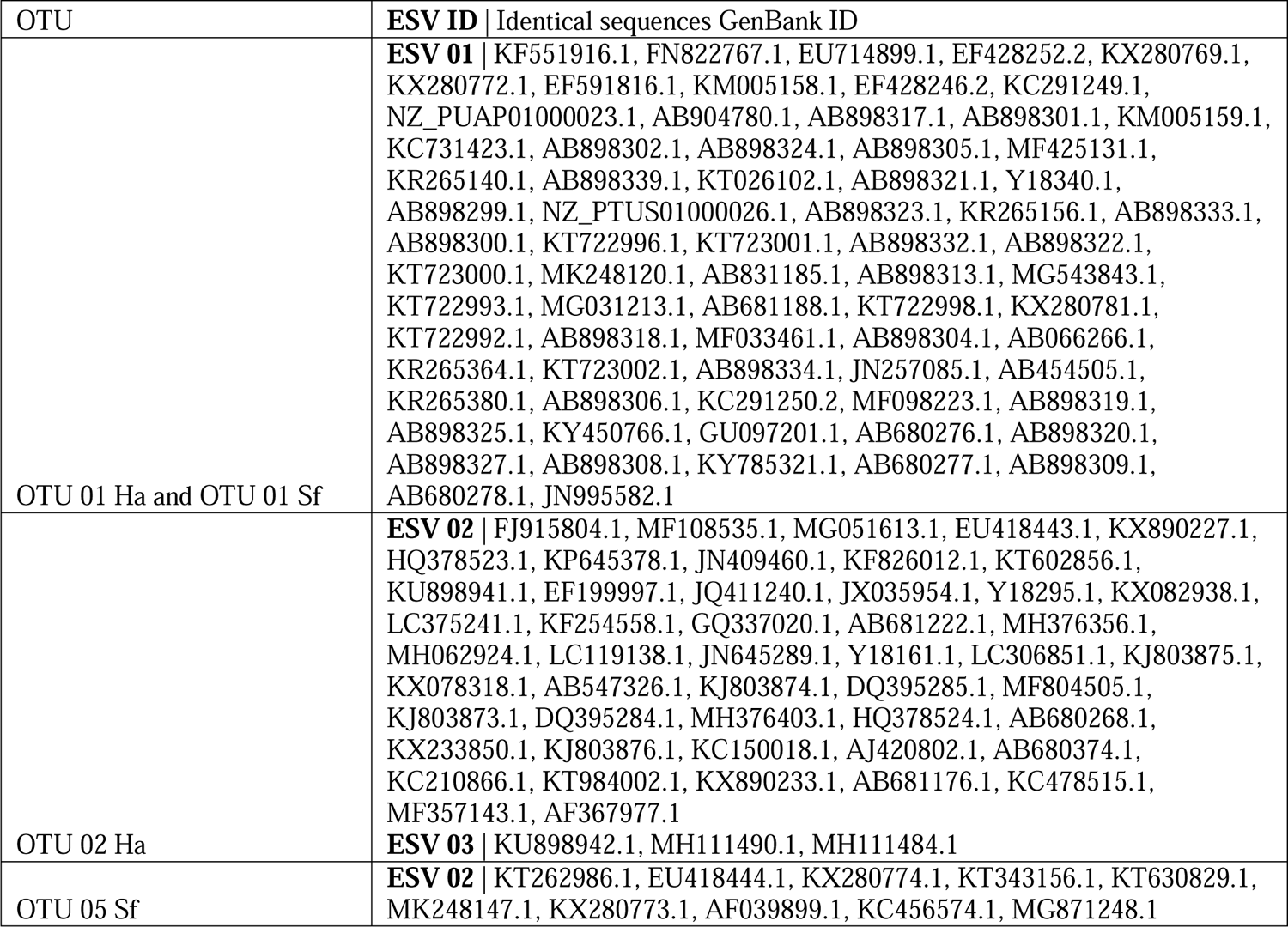
Identical matches in sequence between *Enterococci* found in *Helicoverpa armigera* (Ha) and *Spodoptera frugiperda* (Sf) and sequences used to build the phylogenetic tree (Figure 3).

## Notes

### Competing Interest Statement

The authors have declared no competing interest.

